# The need for high-resolution gut microbiome characterization to design efficient strategies for sustainable aquaculture production

**DOI:** 10.1101/2024.02.29.582783

**Authors:** Shashank Gupta, Arturo Vera-Ponce de León, Miyako Kodama, Matthias Hoetzinger, Cecilie G. Clausen, Louisa Pless, Ana R.A. Verissimo, Bruno Stengel, Virginia Calabuig, Renate Kvingedal, Stanko Skugor, Bjørge Westereng, Thomas Nelson Harvey, Anna Nordborg, Stefan Bertilsson, Morten T. Limborg, Turid Mørkøre, Simen R. Sandve, Phillip B. Pope, Torgeir R. Hvidsten, Sabina Leanti La Rosa

**Affiliations:** Faculty of Chemistry, Biotechnology and Food Science, Norwegian University of Life Sciences, Ås, Norway; Center for Evolutionary Hologenomics, The Globe Institute, University of Copenhagen, Copenhagen, Denmark; Faculty of Biosciences, Norwegian University of Life Sciences, Ås, Norway; Department of Aquatic Sciences and Assessment, Swedish University of Agricultural Sciences, Uppsala, Sweden; Cargill Food Solutions – R&D – SST, Norway; Cargill Aqua Nutrition, Cargill, Sandnes, Norway; Department of Biotechnology and Nanomedicine, SINTEF, Trondheim, Norway; Centre for Microbiome Research, School of Biomedical Sciences, Queensland University of Technology (QUT), Translational Research Institute, Woolloongabba, Queensland, Australia

## Abstract

Microbiome-directed dietary interventions such as microbiota-directed fibers (MDFs) have a proven track record in eliciting responses in beneficial gut microbes and are increasingly being promoted as an effective strategy to improve animal production systems. Here we used initial metataxonomic data on fish gut microbiomes as well as a wealth of a priori mammalian microbiome knowledge on α-MOS and β-mannan-derived MDFs to study effects of such feed supplements in Atlantic salmon (*Salmo salar*) and their hitherto poorly characterized gut microbiomes. Our multi-omic analysis revealed that the investigated MDFs (two α-mannans and an acetylated β-galactoglucomannan), at a dose of 0.2%, had negligible effects on both host gene expression, and gut microbiome structure and function under studied conditions. While a subsequent trial using a higher (4%) dietary inclusion of β-mannan significantly shifted the gut microbiome composition, there were still no biologically relevant effects on salmon metabolism and physiology. Only a single *Burkholderia-Caballeronia-Paraburkholderia* (*BCP*) population demonstrated consistent and significant abundance shifts across both feeding trials, although with no evidence of β-mannan utilization capabilities or changes in gene transcripts for producing metabolites beneficial to the host. In light of these findings, we revisited our omics data to predict and outline novel and potentially beneficial endogenous lactic acid bacteria that should be targeted with future, conceivably more suitable, MDF strategies for salmon.

**IMPORTANCE:** This study focuses on the potential of MDFs to improve aquaculture production. Despite preliminary 16S rRNA amplicon data suggested that populations in the salmon gut microbiome could utilize structurally complex mannans, our findings indicates that endogenous microbes could not metabolize it, nor the host responds to its dietary inclusion, at least not under the trial conditions investigated in this study. We highlight that high-resolution and host-specific microbiome characterization can greatly improve trial design and selection of candidate MDFs for future nutritional interventions. Understanding the intricate interplay between host and its gut microbiome is paramount in studies seeking to leverage endogenous microbial communities to benefit the host. While each new condition, whether it is a disease onset or a nutritional stressor, has the potential to profoundly reshape the microbial diversity, composition and outputs, the functional microbiome information gained under healthy conditions represent a pivotal step towards designing more effective trials involving microbiome-reprogramming feed additives. Overall, we envisage that these results will lead to improved focus on coupling fundamental microbiome characterization to the design of next-generation feeds for salmon aquaculture.

## OBSERVATION

Developing efficient and environmentally sustainable aquaculture production systems is essential to guarantee long-term food security, especially in light of the twofold increase in global demand for seafood expected by 2050 (www.fao.org). Identification of sustainable feed ingredients that promote fish welfare and maximize growth potential, with minimal environmental impacts, is therefore a major research focus in the aquafeed industry. One interesting type of such feed ingredients have little direct nutritional value for the animal, but rather target and shift microbial populations inhabiting the gastrointestinal tract and thereby influence the feed-microbiome-host axis in a beneficial way (1). These microbiome-modulating feed ingredients are well-established in terrestrial animal production (2), but are now also gaining attention in aquafeed research. In salmonid feed research, there have been positive growth effects from supplementing feed with carbohydrates, and for Atlantic salmon fructo-oligosaccharides and α-mannooligosaccharide (α-MOS from yeast cell wall) seems particularly promising (3). In addition, *in vitro* studies with a salmon gut simulator (4) have demonstrated that supplementation of α-MOS leads to shift in microbial community composition, with increase in lactic acid producing *Carnobacterium*, and enhanced production of propanoic and formic acids, both of which have been demonstrated to positively impact animal microbiome and health (5).

Another emerging and potentially powerful feed design strategy is to use microbiota-directed fibres (MDFs). These compounds have chemical structures that align with specific enzymatic capabilities of certain microbial species (6). Examples include β-mannans that are plant-derived glycans found abundantly in human and livestock diets. Depending on their sources, these β-mannans have been categorized into four subtypes; linear β-mannan, galactomannan, glucomannan and galactoglucomannan (6). Norway spruce wood-derived galactoglucomannan, for instance, was developed as an MDF in weaning piglets and was shown to be highly selective for *Roseburia intestinalis* as well as cross-feeding *Faecalibacterium prausnitzii* populations, with directed changes in short chain fatty acid (SCFA) output towards butyrate (6–9).

Despite preliminary indications via taxonomic surveys that *Carnobacterium, Roseburia* and *Faecalibacterium* spp. exist in the salmon gut (10), there is a striking lack of genomic information for the salmon gut microbiome, preventing *in-depth* evaluation of the effectiveness of MDFs such as α- and β-mannans to match enzymatic abilities inherent to endogenous gut microbes. Nonetheless buoyed by the circumstantial evidence surrounding mannans as an MDF, we enthusiastically pursued this uncharted application in a series of trials with the motivation that it could offer exciting prospects for aquaculture research and innovation, potentially aiding salmon feed production to enhance industry sustainability and feed efficiency. We further applied a series of host and microbiome omics analyses with the two-pronged objectives of (i) better understanding the mechanistic link between salmon gut microbes, their metabolic functions, and the host physiology (11, 12) and (ii) evaluating the potential of mannans as MDFs in salmon aquafeed. This included 16S rRNA gene profiling, metatranscriptomics, targeted metabolomics and transcriptomics of salmon organs across feeding trials with different inclusion levels of mannans and across different life stages (**Text S1**.**2**). Our study provides evidence that α- and β-mannan-derived MDFs has negligible effects on salmon, both at the level of gut microbiome and fish physiology. We conclude that these specific MDFs do not qualify for further research as novel feed ingredients in salmon aquafeeds aimed at stimulating microbes present under normal rearing conditions, that is, in the absence of a pathogenic assault or an environmental or nutritional stressor. Nevertheless, the work still showcases the power of our extensive biomolecular data collection along with our newly generated salmon microbial genome atlas (13) to devise promising avenues for testing and using similar feed additives that could have beneficial microbiome-reprogramming outcomes in the salmon gut.

## Results and Discussion

To address knowledge gaps concerning the feed-microbiome-host axis in salmon, a trial was designed to assess individualized responses to an industry-standard 0.2% inclusion of one acetylated β-galactoglucomannan (MN3) and two different types of α-mannans (MC1 and MC2). These MDFs had varying degrees of polymerization and substrate complexity in the form of side-chain decorations and acetylation patterns (**Text S1**.**2**). Samples were collected at different developmental stages of the fish, and varying layers of phenotypic and omics data (16S rRNA, metagenomic, (meta)transcriptomic) was generated from the gut microbiome as well as host gut tissue (**Fig. 1A**). Phenotypic scoring suggested the varying diets had no effect on key performance indicators (KPIs) such as weight, length, and organ integrity (**Fig. 1B, Text S2.1: Table S3**). A series of computational analyses were performed to investigate the correlation between alterations in enzyme produced by diverse microbiota and potential shifts in nutrient utilization or uptake within the fish gut. We detected a total of 839 bacterial genera from 44 phyla using 16S rRNA gene sequencing in a global gut analysis of all fish samples, however no MDF driven structural changes in the gut microbiome were observed (**Fig. 1C, E**), not even as the microbiome evolved over time as the fish transitioned from fresh to saltwater (**Fig. 1D**). An in-depth characterization of microbial functions and community level expression recovered 117,261 microbial genes, and while their activity profiles changed as expected over life stages (**Text S2.3**), but no significant clustering with respect to different MDF (**Fig. 1H**). Further analysis using DESeq2 revealed only 208 significantly differentially expressed genes (DEGs) (ranging from 8-36 between MDFs and control at different life stages) in sampled pre-smolts (T1), smolts (T2) and post-smolts (T3) were identified, none of which were metabolically linked with mannan (**Text S2.3: Table S5**). Finally, to see if diet-driven changes exerted any metabolic influence on their host directly or indirectly (via microbiome activity), we performed transcriptomic analyses on the gut tissue from 48 salmon that were fed either a control diet or the three MDF diets at pre-smolts (T1), smolts (T2) and post-smolts stages (T3). As expected, we found significant differences between life stages (**Text S2.4: Fig. S7**) but did not observe any significant differences in gene expression between the MDF diets and control (**Fig. 1F**).

**FIG 1.**
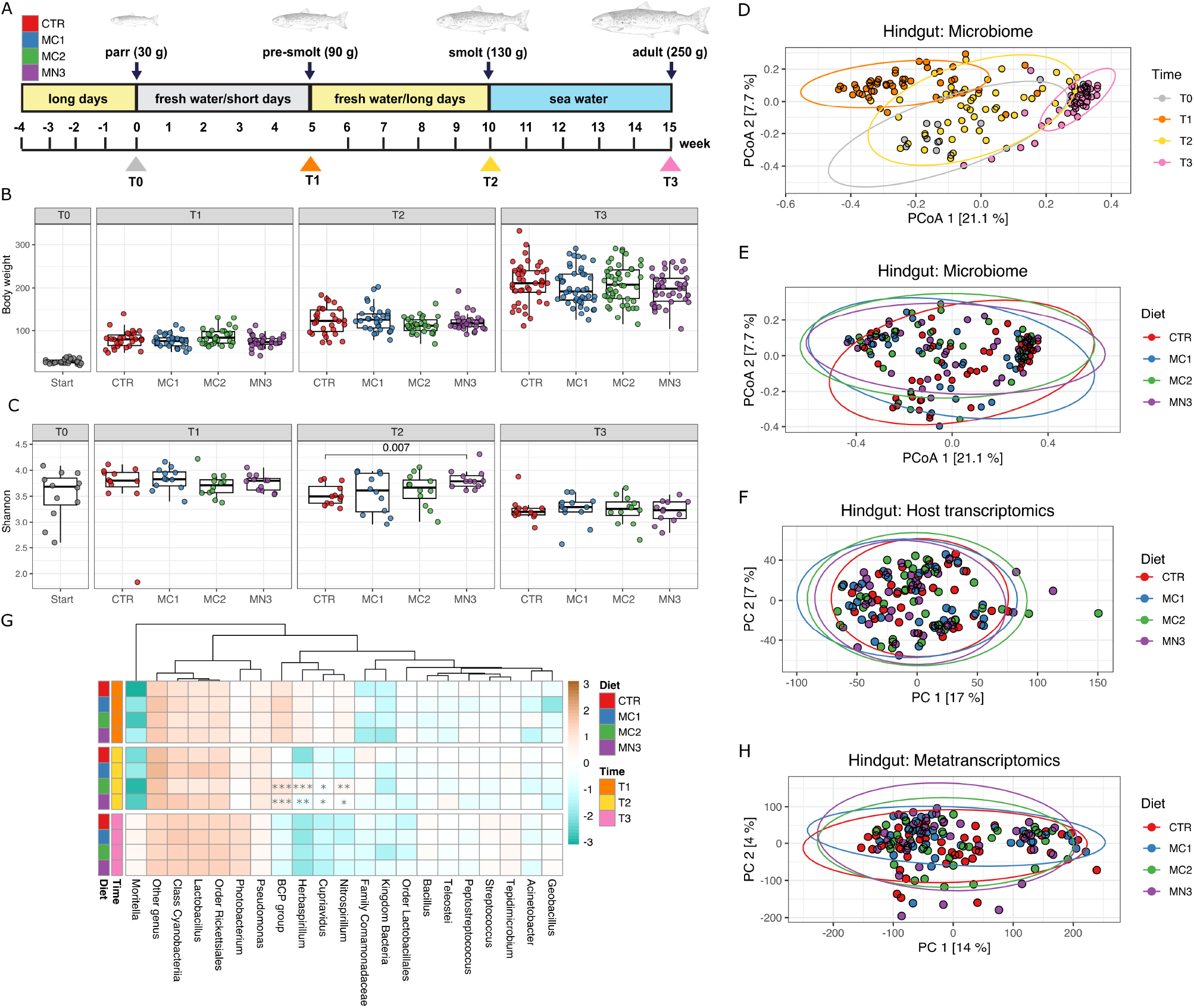
Effect of low dosage of β-mannan on host and gut microbial community structure and function. **(A)** Sampling strategy for studying the effect of low dosage of mannan diet on the temporal dynamics of the Atlantic salmon gut microbiota. T0 (parr), T1 (pre-smolts), T2 (smolts) and T3 (post-smolts) here represents the different sampling time point, and the experimental groups are labeled as CTR (Control), MC1 (Diet 1), MC2 (Diet 2), and MN3 (Diet 3), indicating the different diets administered during the study. **(B)** Box plot showing the mean body weight in all the experimental group, stratified over sampling time. **(C)** Alpha diversity was computed for Shannon and statistically tested using the Wilcoxon test. P-values were corrected using the Benjamin-Hochberg FDR method, with the analysis stratified over time. **(D)** Beta diversity was assessed through Bray-Curtis dissimilarity for 16S rRNA gene hindgut samples, testing the effects of different diets with PERMANOVA. **(E)** Beta diversity was assessed using Bray-Curtis dissimilarity for 16S rRNA gene hindgut samples and assessed over time through PERMANOVA. Each dots represents individual samples colored by different MDFs, as mentioned in the legend. **(F)** PCA plot of gene expression in the hindgut. Plot showing the variance between different MDF and control samples. The percentages on each axis represent the percentages of variation explained by the principal components. **(G)** Top 20 most abundant genera in all the groups. Statistical significance calculated by Wilcoxon test indicated with stars: * p <= 0.05, ** p <= 0.01, *** p <= 0.001. **(H)** PCA plot of metatranscriptome expression in the hindgut. Plot showing the variance between different MDF and control samples. The percentages on each axis represent the percentages of variation explained by the principal components.

To ensure that MDF inclusion levels were high enough to facilitate a diet-driven alteration of the salmon gut microbiome, we performed a small-scale re-iteration of our original trial with freshwater salmon fed a diet supplemented with 4% acetylated galactoglucomannan (**Fig. 2A**), a level that had proven results in monogastric animal trials (7). In this case, the microbiome analysis was extended to include both the hindgut and pyloric caeca, which is a critical part of the salmon digestive system (14, 15). Using a similar data generation and analysis workflow, we identified 683 bacterial genera from 36 different phyla from the hindgut and 510 bacterial genera from 33 phyla from the pyloric caeca samples. The 4% supplementation with galactoglucomannan (4% MN3) in the feed caused a significant change in microbiome composition (Shannon diversity; Wilcoxon, p=0.045, Bray-Curtis distance metrics tested by PERMANOVA for diet, hindgut: p = 0.013, pyloric caeca: p = 0.0035) (**Fig. 2C, D, E and Text S2.5**). Differential genus abundance analysis showed that the *Burkholderia-Caballeronia-Paraburkholderia* (*BCP*) group (Wilcoxon, p<0.001) and *Pseudomonas* (Wilcoxon, hindgut: p<0.001, pyloric caeca: p<0.05) taxa were significantly increased in relative abundance, while levels of *Limosilactobacillus* were reduced (Wilcoxon, p<0.001) (**Fig. 2F**). Despite observing microbiome compositional shifts, no significant changes in host phenotype and metabolism were observed (**Fig. 2B**), with no significant clustering differences in gene expression profiles generated from hindgut **(Fig. 2G)** and pyloric caeca tissues (**Text S2.6: Fig. S9**) of fish fed either the control or experimental diet. A limited number of significant DEGs (only 6 out of 47,563 for hindgut and 2 out of 47,433 genes for pyloric caeca) were identified, none of which were linked to galactoglucomannan degradation (**Text S2.6**). Metatranscriptomic analysis of the hindgut content revealed the expression of 17,094 bacterial genes in both the control group and the high dose galactoglucomannan group, with no significant clustering between them (**Fig. 2H and Text S2.7**). Further analysis using DESeq2 revealed only 5 significant DEGs and none of which were metabolically linked with mannan (**Text S2.7: Table S9**). While targeted metabolomics indicated an increase in acetate concentrations in fish fed 4% β-mannan (t-test, hindgut: p = 1.39e-7, pyloric caeca: p=3.216e-4; **Text S2.8: Table S10**), we observed no metabolic evidence that microbial fermentation pathways linked to its production were significantly increased in gene expression data, nor did we detect endo-mannanases or mannan-specific acetyl esterases (e.g. Carbohydrate Esterase belonging to family 2 and 17) (6, 7), which theoretically could cleave acetyl decorations and depolymerize the galactoglucomannan substrate (**Text S2.5**).

**FIG 2.**
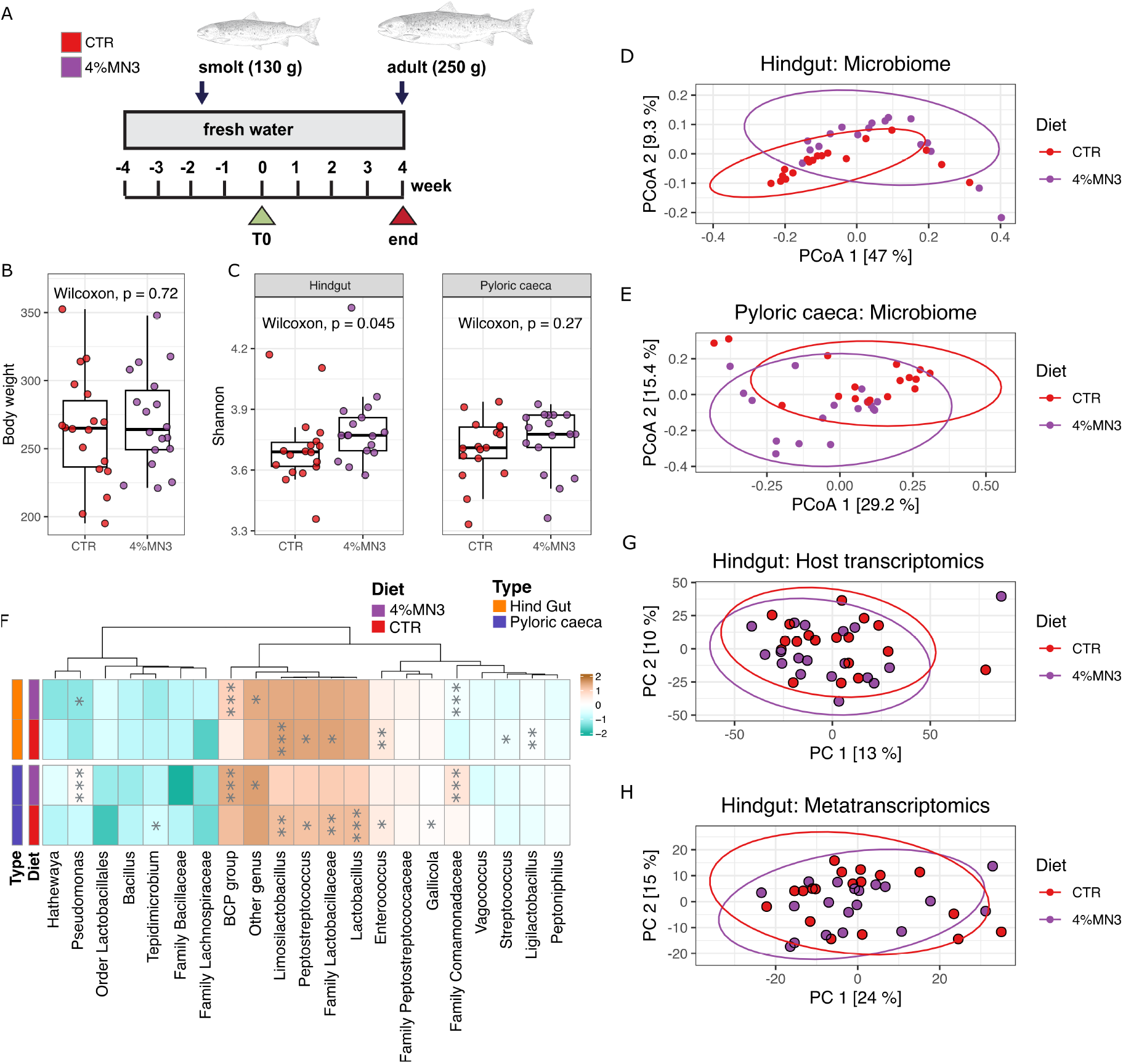
Effect of high dosage of β-mannan on host and gut microbial community structure and function. Sampling strategy for studying the effect of high dosage of β-mannan diet on the temporal dynamics of the Atlantic salmon gut microbiota. The experimental groups are labeled as CTR (Control), and 4%MN3 diet (4% β-mannan diet). **(B)** Box plot showing the mean body weight in all the experimental group. **(C)** Alpha diversity was computed for Shannon and statistically tested using the Wilcoxon test. P-values were corrected using the Benjamin-Hochberg FDR method, with the analysis stratified using sample type that is hindgut and pyloric caeca. **(D)** Beta diversity was assessed through Bray-Curtis dissimilarity for 16S rRNA gene hindgut samples, testing the effects of different diets with PERMANOVA. **(E)** Beta diversity was assessed using Bray-Curtis dissimilarity for 16S rRNA gene pyloric caeca samples and assessed through PERMANOVA. Each dots represents individual samples colored by diet, as detailed in the legend. **(F)** PCA plot of gene expression in the hindgut. Plot showing the variance between MDF and control samples. The percentages on each axis represent the percentages of variation explained by the principal components. **(G)** Top 20 most abundant genera in all the groups. Statistical significance calculated by Wilcoxon test indicated with stars: * p <= 0.05, ** p <= 0.01, *** p <= 0.001. **(H)** PCA plot of metatranscriptome expression in the hindgut. Plot showing the variance between MDF and control samples. The percentages on each axis represent the percentages of variation explained by the principal components.

While we observed a subdued response of microbial and host metabolism to dietary inclusion of mannan, our multi-layered omic analyses still returned valuable information pertaining to microbial functions in the salmon gut microbiome and how they may act to benefit their host. For example, the *BCP group* **(Fig. 1G, Fig. 2F)**, *Limosilactobacillus* and *Lactobacillus* populations all had diet-driven changes in their abundances indicating their potential relationships to external factors that could be leveraged to facilitate their metabolic control (**Fig. 2F, Fig. 3**). Closer examination of their metabolism using genome centric metatranscriptomics highlighted that several enzymatic features and pathways related to pectic galactans, xylans, chitin, beta-glucans, xyloglucans, host mucin-derivatives, celluloses, and undecorated manno-oligosaccharides were putatively active in the salmon gut for the low dose mannan trials (**Fig. 3**). In particular, the lactic acid-producing bacteria (LAB) *Lactobacillus* and *Limosilactobacillus*, which are renowned for their improvements to fish disease resistance via immunostimulation (16), were observed to express genes encoding xylosidases, glucosidases, and galactosidases related to xyloglucans and galactans utilization. These findings conform with their capabilities to metabolize commercially available galacto-oligosaccharides (GOS) previously identified to stimulate growth of similar LAB under *in vitro* culture conditions (17). However, our *in vivo* detection of GH42 β-galactosidases and GH43 arabinofuranosidases point to an improved capacity of using substrates with more complex structure, such as pectin-derived galactans or (arabino)xyloligosaccharides from cereals, than many commercial undecorated GOS (synthesized from monomers obtained from animal products i.e., lactose from milk), and which could be extrapolated into sustainable feed supplements as a LAB-specific MDF (18). We subsequently hypothesize that MDF trials designed around this new microbiome information will result in more concrete insights about the metabolic roles of gut associated microbiota in salmon and other fish in the instances they are implemented.

**FIG 3.**
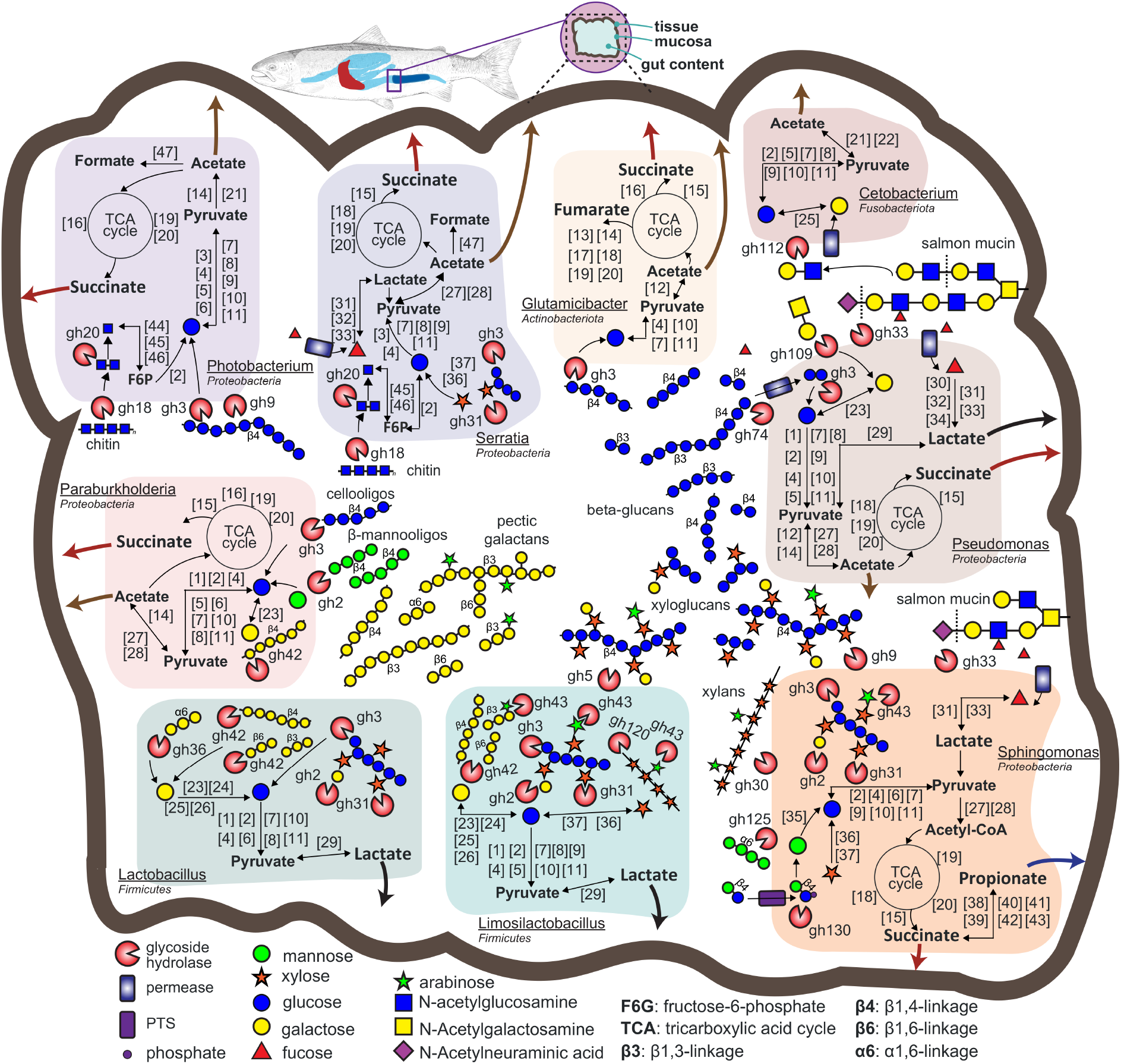
Selected metabolic features of the salmon gut microbiome of adult fish as inferred from genome and metatranscriptome comparisons. The different metabolic pathways (host and dietary carbohydrate depolymerization, glycolysis, TCA and SCFA production are displayed for each population MAG. Graphical representation of different carbohydrates, CAZymes, and cellular features are based on functional annotations that are depicted as numbered or abbreviated gene boxes, which are additionally listed in **Table S4**. Features are included if a gene was expressed at either the smolt (T2) or adult (T3) stage from either the control or MDF (MC1, MC2, MN3) diets. The main carbohydrates (betaglucans, xylans, galactans, and chitin) predicted to be utilized, SCFAs (i.e., acetate), and organic acids (like lactate and succinate) are represented by large colored arrows. GTBD-Tk inferred taxonomy is included. Gene names and abbreviations are also provided in **Table S4**.

In conclusion, whilst confronted with promising preliminary 16S rRNA sequencing data suggesting that mannans would selectively target beneficial populations allegedly inherent to the salmon gut microbiome, our findings instead showed that their dietary inclusion had negligible impact on microbial functionalities and host physiology and metabolism. This was largely demonstrated by in depth analyses of trial KPIs, microbial (meta)genomes as well as both host and microbial RNA and metabolites, which highlighted a scarcity of mannan-degrading capabilities but in its place recovered in-depth, first-time *in vivo* characterization of populations that in retrospect should have acted as targets for MDF design and application. Our experiences highlight the risk of inferring functional outcomes from 16S amplicon data and clearly reinforce the impact that critical metabolic insight can have on microbiome intervention strategies, strengthening the need for high-resolution, host-specific, microbiome characterization as a prerequisite to improved animal trial design. Further, our results also highlight the value of focusing on microbes that are naturally present in the host species of interest and the metabolic capacity of these microbes towards identifying the next pre- and probiotic candidates. While establishing a baseline understanding of the host’s microbial community under normal healthy conditions, the next crucial step is to characterize changes in microbiota in altered states, such as under environmental stressors or in a disease state, with the goal to identify and broaden potential targets for intervention by MDFs. Criticisms towards metataxonomy-based microbiome characterization as being overly descriptive or merely stamp-collecting are easy to make, but the value of microbial genome atlases and culture biobanks (13) cannot be exaggerated, especially as industry makes increasingly strong moves towards precision microbiome interventions as a viable technology to improve aquaculture sustainability and production.

## Data availability

To access the code and data used in this study, please refer to our GitHub repository (https://github.com/shashank-KU/ImprovaFish-MDF-Effects). The raw metagenomics dataset analyzed during the current study for low and high dose trials will be available, upon publication, in the Sequence Read Archive (SRA) repository under project id PRJNA947090. The raw host transcriptomics and metatranscriptomics data for the low dose mannan trial are available under project id PRJEB73366 and PRJEB67787, and for the high dose mannan trial under project id PRJNA1051365 and PRJNA1051380, respectively.

## ACKNOWLEDGMENTS

This work was supported by the Research Council of Norway (project no. 300846), the Swedish Research Council Formas (grant no. 2019-02336) and the European Union’s Horizon 2020 research and innovation program under the ERA-Net Cofund project BlueBio (grant agreement no. 311913). The Danish National, Research Foundation grant no. DNRF143 to M.T.L. The Orion High Performance Computing Center at the Norwegian University of Life Sciences and Sigma2 - the National Infrastructure for High Performance Computing and Data Storage in Norway are acknowledged for providing computational resources that have contributed to meta-omics analyses described in this study. Jacob A. Rasmussen is thanked for help handling and depositing raw sequence data. The authors thank Elixir-Norway (NFR project no. 322392) for bioinformatics and data management related services.

## AUTHOR CONTRIBUTIONS

P.B.P., S.L.L.R., S.R.S., S.B and T.R.H. designed the study. Transcriptomics and meta-transcriptomics data generation were done by M.T.L, C.G.C., L.P., and A.R.A.V. Transcriptomics and meta-transcriptomics analysis were done by S.G., and M.K. 16S rRNA gene sequencing analysis were carried out by S.G. Metabolomic analyses were carried out by S.G., and S.L.L.R. T.N.H., and M.H. helped in the sequence analysis. The draft manuscript was written by S.G., S.L.L.R., P.B.P, S.R.S., and T.R.H. All authors contributed to the editing of the text and content and approved the final version.

We declare no competing financial interests.

